# Sustained pigmentation causes DNA damage and invokes translesion polymerase Pol κ for repair in melanocytes

**DOI:** 10.1101/2022.05.20.492761

**Authors:** Shivangi Khanna, Madeeha Ghazi, Yogaspoorthi Subramanian, Farina Sultan, Iti Gupta, Kanupriya Sharma, Sudhir Chandna, Rajesh S Gokhale, Vivek T Natarajan

## Abstract

The pigment melanin protects skin cells from ultraviolet (UV) radiation induced DNA damage. However, intermediates of eumelanin are highly reactive quinones that are potentially genotoxic. In this study, we systematically investigate the effect of sustained elevation of melanogenesis and map the consequent cellular repair response of melanocytes. Pigmentation increases DNA damage, causes cell cycle arrest, and invokes translesion polymerase Pol κ for DNA repair in primary human melanocytes, as well as mouse melanoma cells. Confirming the causal link, CRISPR-based genetic ablation of tyrosinase, the key melanin synthesizing enzyme results in depigmented cells with low Pol κ levels. However, silencing of Pol κ in pigmenting cells results in unchecked proliferation despite the presence of damaged DNA, that could potentially lead to genome instability. Thereby, our results indicate Pol κ to be a necessary evil to resolve melanin induced damage. Error-prone repair by Pol κ in part explains the mutational landscape observed in human melanoma. Thus, our study illuminates a hitherto unknown dark side of melanin and identifies (eu)melanogenesis as a key missing link between tanning response and mutagenesis mediated *via* the Pol κ-based low fidelity DNA repair response of melanocytes.

**Key Highlights:**

- Sustained melanogenesis causes DNA damage in melanocytes
- Melanogenesis elicits replication stress and translesion repair by Pol κ
- Pol κ resolves melanin-induced DNA damage and suppresses genome instability
- Expression of Pol κ correlates with mutational load in human melanoma

## Introduction

Skin pigmentation acts as an important barrier against penetration of harmful ultraviolet (UV) radiations. Melanin polymer with its broad absorption spectrum protects the genome from damage by high energy UV rays (Brenner & Hearing, 2008). Hence skin tanning is an essential physiological response for protection against UV-induced mutagenesis and consequent risk of malignancy. The adaptive nature of pigmentation is evident in world-wide distribution of basal skin tone as well as the extent of tanning in populations residing across latitudes that differ in incident UV flux (Yamaguchi et al., 2007). The presence of melanin laden melanosomes in keratinocytes protects these cells from UV damage (Ferreira et al., 2013). However, synthesis of melanin within melanocytes poses challenge as the intermediates are highly reactive quinones and potentially genotoxic genotoxic (Miranda et al., 1994; Pawelek & Lerner, 1978). Variety of factors contribute to the mutation burden in melanocytes, among these the role of melanogenesis, if any remains obscure.

The nature of DNA damage caused by UV-A, UV-B and UV-C, as well as the cellular response are very well-characterized (Anna et al., 2007; Rastogi et al., 2010). UV-signature mutations arise in keratinocytes consequent to the repair of DNA photoproducts, such as cyclobutane pyrimidine dimers (CPDs) and pyrimidine 6-4 pyrimidones (6-4 PP). These are resolved via Nucleotide Excision Repair (NER) pathway. The autosomal recessive syndrome Xeroderma Pigmentosum caused by mutations in various genes in this pathway and the translesion polymerase eta (PolH) render individuals sensitive to sunlight (Masutani et al., 1999) (D’Errico et al., 2003). Tumor suppressor p53 is the key orchestrator of UV-mediated DNA damage in keratinocytes, that induces POMC gene expression, resulting in α-MSH secretion (Cui et al., 2007). In neighboring melanocytes, α-MSH activates the downstream MC1R signaling. In addition to promoting melanogenic response, this pathway invokes NER which pre-emptively prepares melanocytes for DNA repair (Jarrett et al., 2014; Swope & Abdel-Malek, 2018).

Genetic risk factor of melanoma links to the red hair phenotype of individuals with lighter skin tone due to the presence of pheomelanin. Systematic studies have elucidated the role of oxidative damage in UV-independent carcinogenesis, induced by pheomelanin (Mitra et al., 2012). Epidemiological studies suggest that the highest melanoma risk is associated with intermittent exposure to UV rays (Armstrong & Cust, 2017). UV-induced free radicals excite electrons in melanin to create a quantum triplet state that transfer energy to DNA and initiate characteristic UV-signature C→T transitions (Premi et al., 2015). Hence, UV directly and in conjunction with melanin is likely to cause DNA damage and initiate mutations. Additionally, as pheomelanin is capable of mutagenesis independent of UV, it is generally believed that formation of eumelanin is protective and hence its role in causing DNA damage is not systematically investigated. In a recent work, investigators identified that melanocytes contain genomic locations that are prone to modifications in several loci implicated in melanoma (Premi et al., 2019). We hypothesized that melanin intermediates could modify DNA and elicit counteracting cellular response.

To investigate this, in the current study we employ a synchronized, temporally resolved pigmenting system using B16 melanoma cells that produce only eumelanin and confirm our results in primary human melanocytes. In a UV-independent manner, the rate of eumelanin production is further enhanced with substrate tyrosine, suppressed with chemical inhibition using Phenylthiourea (PTU) or CRISPR-based mutagenesis. Prolonged elevation of eumelanin synthesis causes DNA damage and melanocyte elicits repair *via* translesion polymerase K (Pol κ) through ATR-Chk1 signalling. In the absence of Pol κ, sustained melanogenesis results in continued proliferation of melanocytes with damaged DNA, indicating that absence of Pol κ could lead to genome instability. Meta-analysis of TCGA-melanoma data shows elevated mutation burden in human patient samples with high *POLK* mRNA levels. Our study thereby establishes the DNA damaging effects of eumelanogenesis and highlights the counteracting role of DNA repair by Pol κ.

## Results

### Melanogenesis induces generation of free radicals and delays cell cycle progression

To systematically investigate the effect of melanogenesis on DNA damage we resorted to the use of B16 pigmentation system that permits the kinetic study of pigmentation and associated cellular changes in melanocytes (Natarajan et al., 2014). Choice of B16 cells that synthesize purely eumelanin, further enabled segregating the role of pheomelanin (Ito et al., 1988). To further perturb the pigmentation, in this study we compare the basal pigmentation state with hyperpigmentation induced by the treatment of 1mM L-Tyrosine (Tyr) that serves as a substrate for tyrosinase enzyme and augments melanogenesis (**Fig S1A**). By the pharmacological inhibition of Tyrosinase with 200 µM Phenylthiourea (PTU), we achieve decreased pigmentation in this progressive pigmentation model (**Fig 1A**).

**Figure 1:**
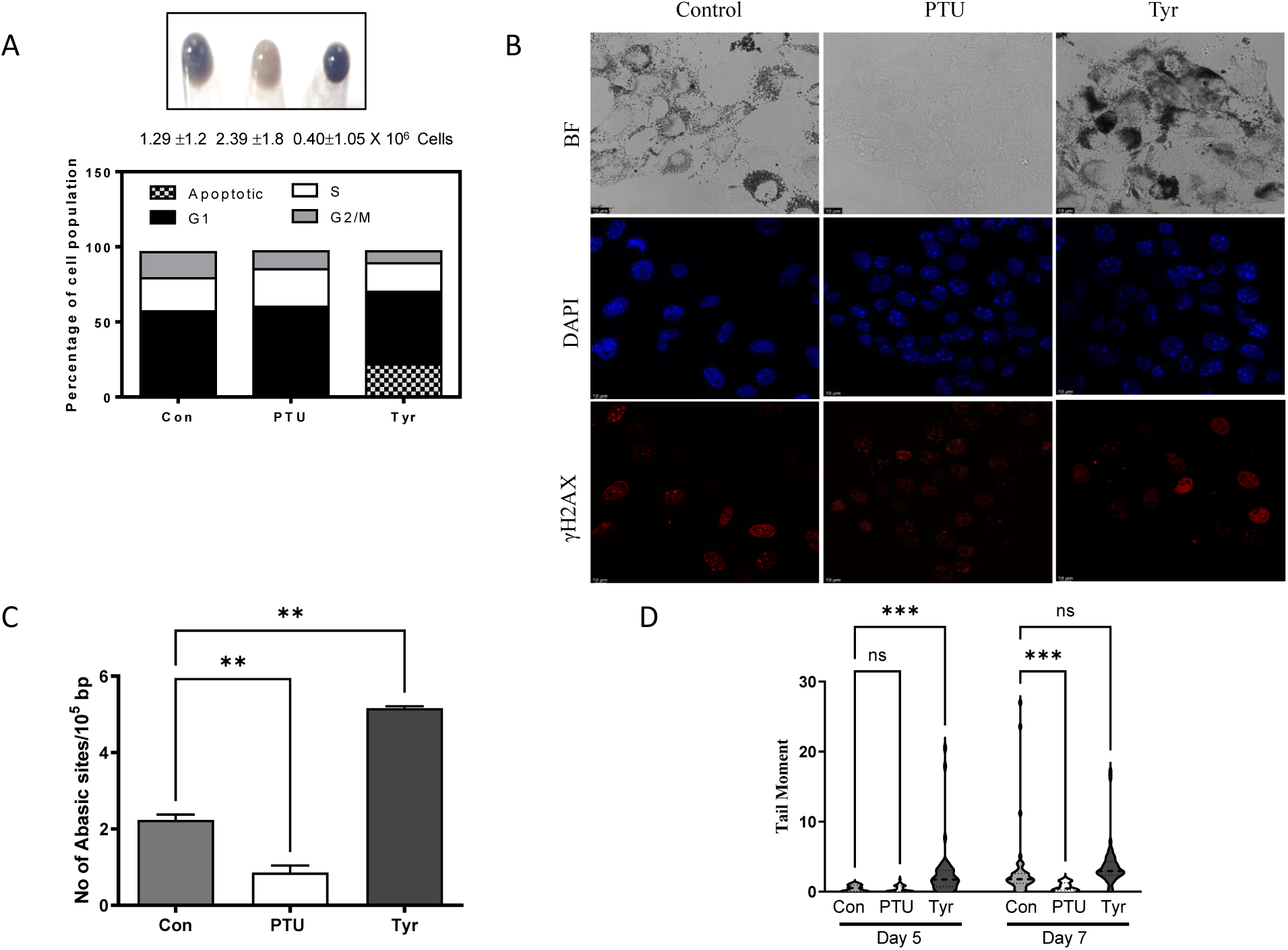
Melanin synthesis causes DNA damage. A. (Top) Cell pellet of B16 mouse melanoma cells grown at low density. Cells were left untreated for control or treatment with tyrosinase inhibitor 200 µM phenylthiourea (PTU) or a 1mM tyrosinase substrate L-Tyrosine (TYR) for seven days. Number of cells, mean ± SEM across biological replicates is depicted below. (Bottom) Cell cycle analysis of cells carried out using Annexin V and propidium iodide staining detected by FACS is depicted as stacked bars. Sub G0 population is marked as apoptotic. Experiment was carried out in triplicates across biological replicates and a representative plot is depicted. B. Immunofluorescence of B16 cells treated with PTU and Tyr with phosphorylated H2AX antibody that labels double strand DNA breaks. C. Number of abasic sites in the genomic DNA was estimated by an aldehyde specific conjugation of biotin and subsequent detection using streptavidin based enzymatic detection. Using standard probes number of abasic sites per 10^5^ bp is estimated. Bars represent mean ±SEM across duplicate biological experiments. students unpaired t test * p val< 0.02 ** p val < 0.002 D. Single cell electrophoresis followed by comet analysis of B16 cells undergoing varying levels of pigmentation in the presence of PTU and Tyr. Experiment was carried out at mid phase (day 5) and late phase (day 7) of pigmentation. Mean tail moment distribution across each population of duplicate biological experiments with atleast 50 comets analyzed is depicted by a violin plot. Error bars represent ± SEM. students unpaired t test ns non-significant, ** p val< 0.01 **** p val < 0.0001

We observed a consistent reduction in the number of cells at day 7 upon hyper-pigmentation with Tyr and an increase upon depigmentation with PTU, compared to control moderately pigmented cells (**Fig 1A**). Cell cycle analysis using propidium iodide staining followed by flow cytometry analysis to quantify cells in different phases of the cell cycle revealed that hyperpigmented cells have reduced S+G2/M population suggesting a reduction in proliferation. Additionally, we observed a small but significant proportion of cells with sub G0 DNA content, indicative of cell death. Staining of differentially pigmented cells at day 7 with annexin V and propidium iodide revealed that hyperpigmented cells have a significantly higher proportion of early and late apoptotic cells (**Fig S1B**). Treatment with PTU reduced these populations, suggesting that uncontrolled hyperpigmentation could result in cell death. Treatment of Tyr and PTU did not alter the cell cycle profile of the B16 cells in high density, non-pigmentation permissive conditions, confirming the effect to be specific to pigmentation (**Fig S1C**). This was further validated by the reduction in S+G2/M population as the melanin progressively accumulates during different days of pigmentation (**Fig S1D**).

### Melanogenesis causes DNA damage

Melanin is a stable free radical, however its synthesis results in the generation of reactive quinone intermediates that can pass the limiting melanosome membrane and result in damage (Riley, 1988). By cytometric DCFDA staining of differentially pigmented cells we observed the highest labelling in Tyr treated, hyperpigmented cells and least in PTU treated cells in which melanin synthesis is minimal (**Fig 1B** & **Fig S1E**). We surmised that these free radicals could cause cellular DNA damage, which could contribute to the observed cell death in hyperpigmented cells. Substantiating this notion, the content of abasic sites in the genomic DNA followed the same pattern as cellular free radical levels (**Fig 1C**). However, DNA base oxidation as detected by 8-oxo-deoxy guanosine levels were comparable across the three pigmentation groups (**Fig S1F**).

Therefore, we hypothesized that melanogenesis could cause DNA damage. Pattern of nuclear γH2AX foci that is associated with DNA strand breaks due to damage, confirmed that the hyperpigmented condition has elevated number of puncta and pretreatment with the melanogenesis inhibitor PTU, reduces the foci formation (**Fig 1B**). Surprisingly not only the γH2AX phosphorylation were elevated in Tyr and decreased in PTU, the total H2AX levels also followed a similar trend (**Fig S2A**). Damage in DNA was further confirmed by assessing the DNA breaks using comet analysis. As the pigmentation progressed, the level of associated DNA nicks increased from day 5 to day 7, and Tyr treated hyperpigmented cells demonstrated maximal DNA nicks and PTU treated depigmented cells had the least amount (**Fig 1D** & **Fig S2B**). In all, these experiments suggest that ongoing melanogenesis could cause damage to the DNA. Comet is also indicative of ongoing repair to reverse the damage, which would transiently increase the number of nicks in DNA (Azqueta et al., 2014; Chandna, 2004). Hence it is likely that ongoing melanogenesis intermediates could cause damage to DNA, and consequently elicit appropriate machinery for repair.

### Melanin induced DNA damage elicits replication stress response

To identify the cellular machinery that could repair melanin induced DNA damage, we resorted to mapping the damage induced changes in DNA repair genes. Based on our earlier work we had profiled the progressive stages of pigmentation of B16 melanocytes (Natarajan et al., 2014). Analysis of genes related to DNA repair across different stages of pigmentation, revealed an interesting pattern. DNA repair genes were elevated, and DNA replication-coupled repair genes were downregulated with progressive pigmentation (**Fig S2C**). To decipher the key component that is involved in repair of melanin-induced DNA damage, we compared the response of B16 melanocytes with defined DNA damaging agents. Towards this we treated either unpigmented or pigmented day 7 B16 cells with UVA or UVB to assess photo-oxidative damage, hydroxyurea for replication stress and H_2_O_2_ for oxidative damage. Pigmented and unpigmented cells that were not treated with any DNA damaging agents were included, and expression of a panel of DNA repair genes were analyzed by qRT-PCR analysis after 24h treatment. The overall response of pigmented and unpigmented cells to various damages were comparable with some prominent differences such as the upregulation of BRCA1 gene to all challenges in pigmented but not in depigmented cells. The signature pattern of expression for each of the damage was clearly distinct, and notably the pattern of gene expression induced by pigmentation closely paralleled that of hydroxyurea treatment that results in replicative stress response (**Fig 2A**).

**Figure 2:**
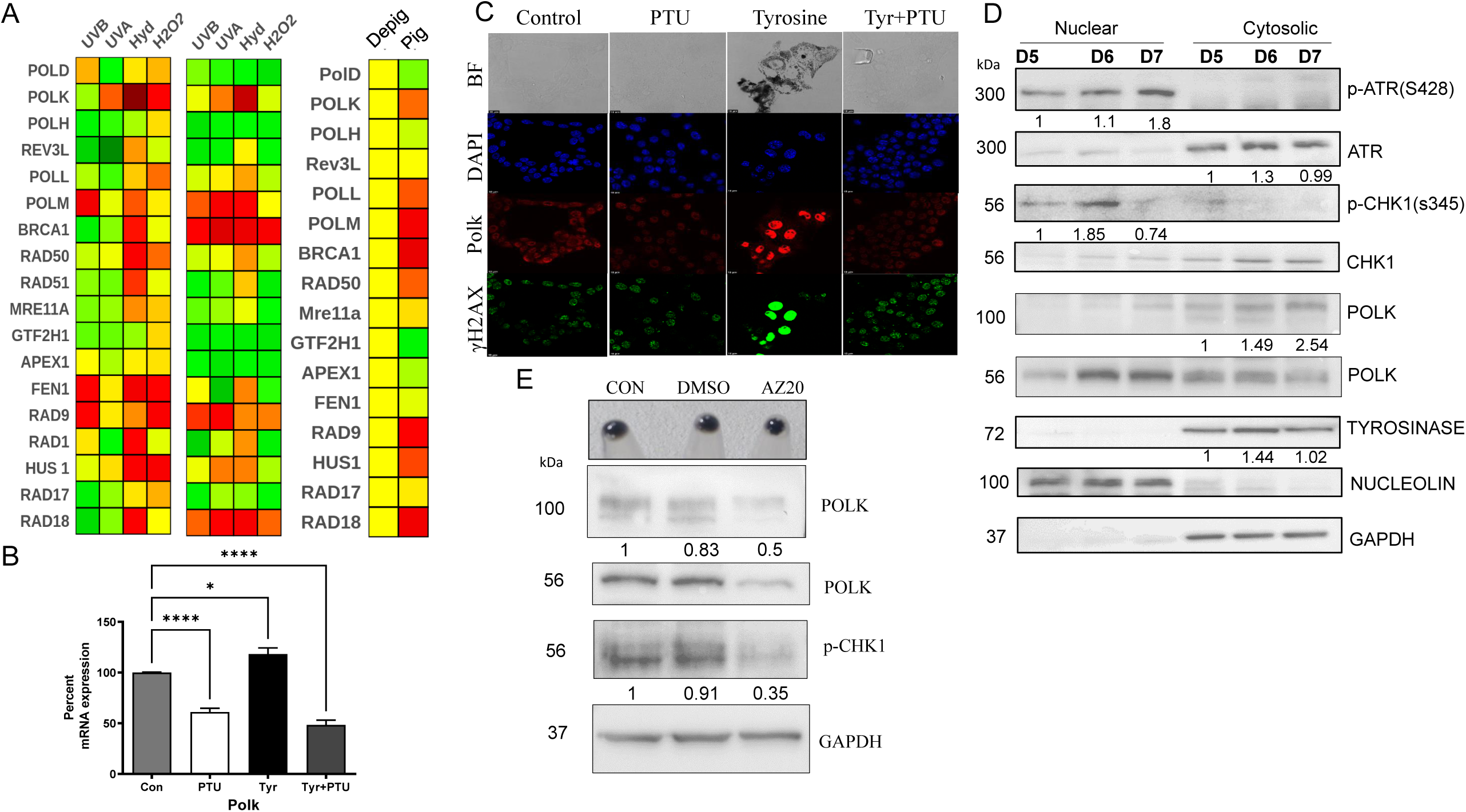
Melanogenesis induces replicative stress response and induces translesion polymerase *Polk*. A. Fold change in mRNA levels of a panel of DNA repair genes by qRT-PCR analysis. The heat map represents fold change and compares DNA repair gene signature of depigmented (left), pigmented (middle) B16 cells with various DNA damaging agents. Untreated pigmented cells compared to depigmented control is depicted as a heat map (right). B. mRNA levels of *Polk* in B16 cells that are allowed to pigment differentially in the presence of 200 µM PTU, 1mM Tyr or both. Bars represent percent mRNA levels compared to control as mean ± SEM across five biological replicates. C. Immunofluorescence of B16 cells treated with 200 µM PTU, 1mM Tyr or both. Top panel is the bright field and dark granules are the pigment accumulation. Nuclear DNA stained with DAPI (blue) and Pol κ in red. D. Western blot analysis of nuclear and the post-nuclear cytoplasmic lysates of B16 cells on day 5 (early), day 6 (mid) and day 7 (late) stages of pigment accumulation. Numbers below the blot correspond to Day 5 normalized expression of the indicated protein. E. Western blot analysis of B16 cells treated with DMSO or 50 nM AZ20 a selective inhibitor of ATR kinase. Numbers below the blot correspond to control normalized expression of the indicated protein wrt *Gapdh*.

### In response to melanogenesis, translesion polymerase Pol κ is induced by ATR-Chk1 pathway

Kinetic profiling of expression changes in DNA repair genes on different days of progressive pigmentation revealed that several homologous recombination repair genes were increased in the expression at early and mid-phase of pigmentation (**Fig S3A**). Interestingly H2AX mRNA followed similar pattern and is in concordance with the observed changes in total H2AX levels by western blot analysis. Strikingly among the translesion polymerases, Pol κ showed a steady increase along with pigmentation at both RNA and protein levels and emerged as a prominent candidate (**Fig S3B-D**). To investigate whether its regulation is critically dependent on pigmentation, B16 cells were treated with PTU, Tyr or both and on day 7 were subject to qRT-PCR analysis for *Polk*. Regulation of *Polk* paralleled the observed pigmentation (**Fig 2B**). Whereas, other two translesion polymerases *Polh* and *Rev3l* were minimally altered and did not show discernable trend in their regulation with pigmentation **(Fig S3E** & **F**). Immunocytochemical localization of Pol κ and labelling of proliferation by EdU (Ethynyl deoxyuridine) was simultaneously carried out in differentially pigmenting B16 cells. We observed high Pol κ as well as γH2AX labelling in tyrosine treated hyperpigmented cells (**Fig 2C**). Prevention of pigmentation by PTU reversed the levels of both Pol κ and γH2AX staining substantiating the induction to be pigmentation dependent. Cell fractionation followed by western blot analysis on different days of pigmentation confirmed induction of Pol κ in the progressive pigmentation model (**Fig 2D**). From these lysates, a clear induction of nuclear localized phosphorylated ATR and phosphorylated Chk1 further indicated activation of DNA repair response pathway during progressive pigment induction.

Having observed an induction of Pol κ during pigmentation and a concomitant increase in the ATR-Chk1 signaling axis, we hypothesized that the ATR-Chk1 pathway could be responsible for Pol κ induction. To test this, during pigmentation, we treated the cells with 50 nM of ATR inhibitor AZ20. Western blot analysis confirmed decrease in p-Chk1 levels and reduction in both 100 kDa and 56 kDa forms of Pol κ (**Fig 2E**). Thereby, using chemical modulators that work on the rate limiting enzyme tyrosinase to modulate pigmentation, we identify Pol κ induction to be under the control of ATR-Chk1 signaling axis. While all these pharmacological approaches pointed towards pigmentation dependent induction of Pol κ, to unequivocally establish the dependence, we resorted to a genetic method of altering the pigment content.

### Genetic ablation of melanogenesis curtails DNA damage and repair response

Though PTU and Tyr did not alter the DNA damage of pigmentation incompetent cells, it is likely that the molecules could trigger the changes in Pol κ levels in cells independent of pigmentation. Hence, we genetically ablated Tyrosinase, the key enzyme involved in melanin synthesis using CRISPR based methodology (**Fig S4A**). The strategy involved transfection of high density B16 cells with synthesized guide RNA (sgRNA) and Cas9 protein complex and plating them at a low density to initiate pigmentation. The depigmented colonies were trypsinized and screened for sequence alterations in the Tyr gene. The identified mutation results in a single base deletion downstream to the sgRNA sequence (**Fig S4B**), resulting in a frameshift mutation (Tyr^fs118^) encoding a truncated protein (**Fig S4C**). Western blot analysis of lysates from different days of pigmentation from wild-type and Tyr^fs118^ indicated a lower level of protein expression due to heterozygous nature of the mutation. Interestingly, during increased pigmentation from day 5 to day 7 the level of TYR and related DCT proteins do not increase appreciably in the mutant in concordance with the interdependent stability of melanosomal proteins and explains the decreased pigmentation observed (**Fig S4D** & **Fig 3A** top panel). Higher proliferation lower DNA damage was observed when the mutant cells were subjected to pro-pigmenting conditions (**Fig S4 E-G**).

**Figure 3:**
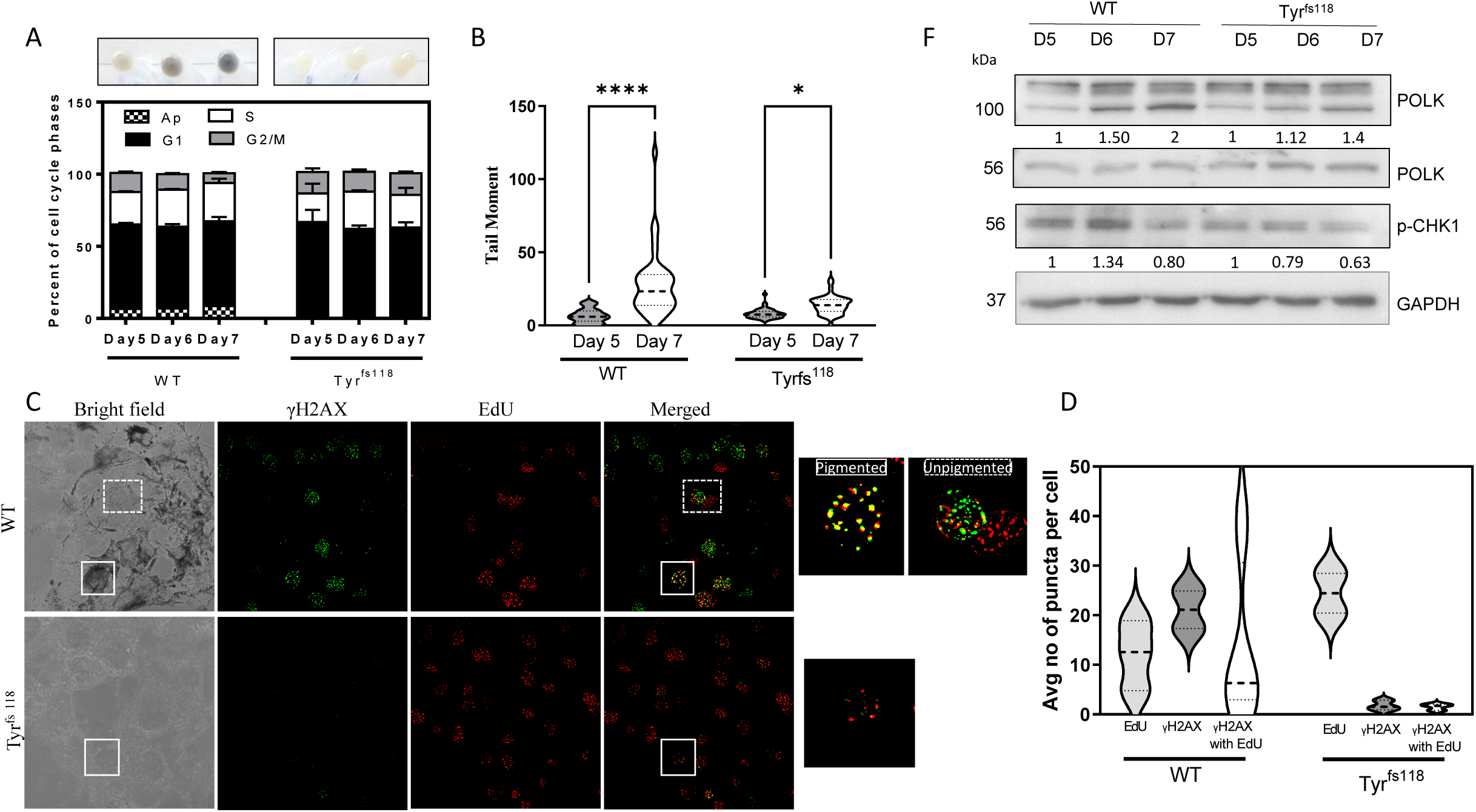
Genetic ablation of tyrosinase by CRISPR prevents melanogenesis, reduces DNA damage and curtails Polk induction. A. Cell pellets of B16 WT and Tyrosinase mutant (Tyr^fs118^) cells on days 5, 6 and 7 of pigmentation (top). Cell cycle analysis by propidium iodide staining, sub G0 cells are labelled apoptotic (below). The stack bars represent mean ± SEM across duplicate experiments. B. B16 WT and Tyr^fs118^ cells on days 5 and 7 of pigmentation were subjected to single cell electrophoresis and comet analysis was performed. Mean tail moment distribution across each population of duplicate biological experiments with atleast 50 comets analyzed is depicted by a violin plot. Error bars represent ± SEM. **** p val < 0.0001 C. Confocal images of B16 WT and Tyrosinase mutant (Tyr^fs118^) that were allowed to pigment for 7 days. Bright field images indicate the presence of melanin granules. Immunofluorescence using EdU (red) and γH2AX antibody (green) is labelled. Merged images show co-localization of γH2AX with EdU (yellow). A single cell showing co-localization is shown as an inset. D. Quantitation of mean of the number of γH2AX puncta that are single positive or double positive (colocalizing with EdU puncta) per cell across 50 cells in B16 WT and Tyrosinase mutant (Tyr^fs118^) cells is depicted stacked bars representing mean ± SEM.

Cell cycle analysis of progressive pigmentation of B16 WT and B16 Tyr^fs118^ cells suggested that along with the accumulating pigmentation the wild type cells demonstrated a progressive decrease in S+G2/M population and an increase in dead cell count. The cell cycle profile of the mutant remained unaltered and even on day 7 sub G0 dead cells were minimal (**Fig 3A**). Associated DNA breaks determined by comet analysis indicated a similar trend and the mutant depigmented cells had minimal comets (**Fig 3B**). These two observations were corroborated by the double staining with γH2AX for nuclear damage/repair foci and detection of newly synthesized DNA using EdU coupled fluorophore labelling (**Fig 3C&D**). While the DNA damage was higher in pigmented cells with a lower proliferation, the mutant cells were depigmented, proliferated more, and had minimal DNA damage. Hence establishing pigmentation to be a significant source of DNA damage in melanocytes. In the WT pigmented cells there was a significant overlap of the two signals indicating that these are regions of damage where new DNA synthesis is ongoing. Interestingly, the wild type cells showed heterogeneity in double stained population. On closer analysis, the heavily pigmented cells had several double positive foci, whereas cells with moderate pigmentation had lower levels of colocalized puncta. Analysis of the EdU and γH2AX colocalized spots clearly suggested that the pigmented cells had a higher proportion of damaged DNA undergoing repair, that could possibly be attributed to translesion synthesis.

### Pigmentation induced DNA damage and elicitation of Pol κ response is recapitulated in Primary Human Epidermal Melanocytes

Since B16 cells are derived from mouse melanoma, it is likely that the observed response could be restricted to melanoma cells and not a physiological response of melanocytes. Hence, we subjected Normal Human Epidermal Melanocytes (NHEM) that are derived from healthy skin and represent the primary melanocytes to differential pigmentation with PTU and Tyr for seven days (**Fig 4A** top panel). Assessment of H2AX phosphorylation by western blot analysis revealed a similar pattern of increased phosphorylation and a concomitant increase in total protein level with hyperpigmentation (**Fig 4A** bottom panel). This was recapitulated in the foci formation detected by immunofluorescence analysis (**Fig 4B**). Comet assay revealed that the DNA breaks were highest in hyperpigmented cells and lowest in depigmented cells (**Fig 4C**). Hence, these observations indicate that pigmentation is indeed a strong physiological response that causes significant DNA damage in non-transformed primary cells. We therefore propose that biochemical intermediates of melanogenesis are potentially genotoxic and cause DNA damage.

**Figure 4:**
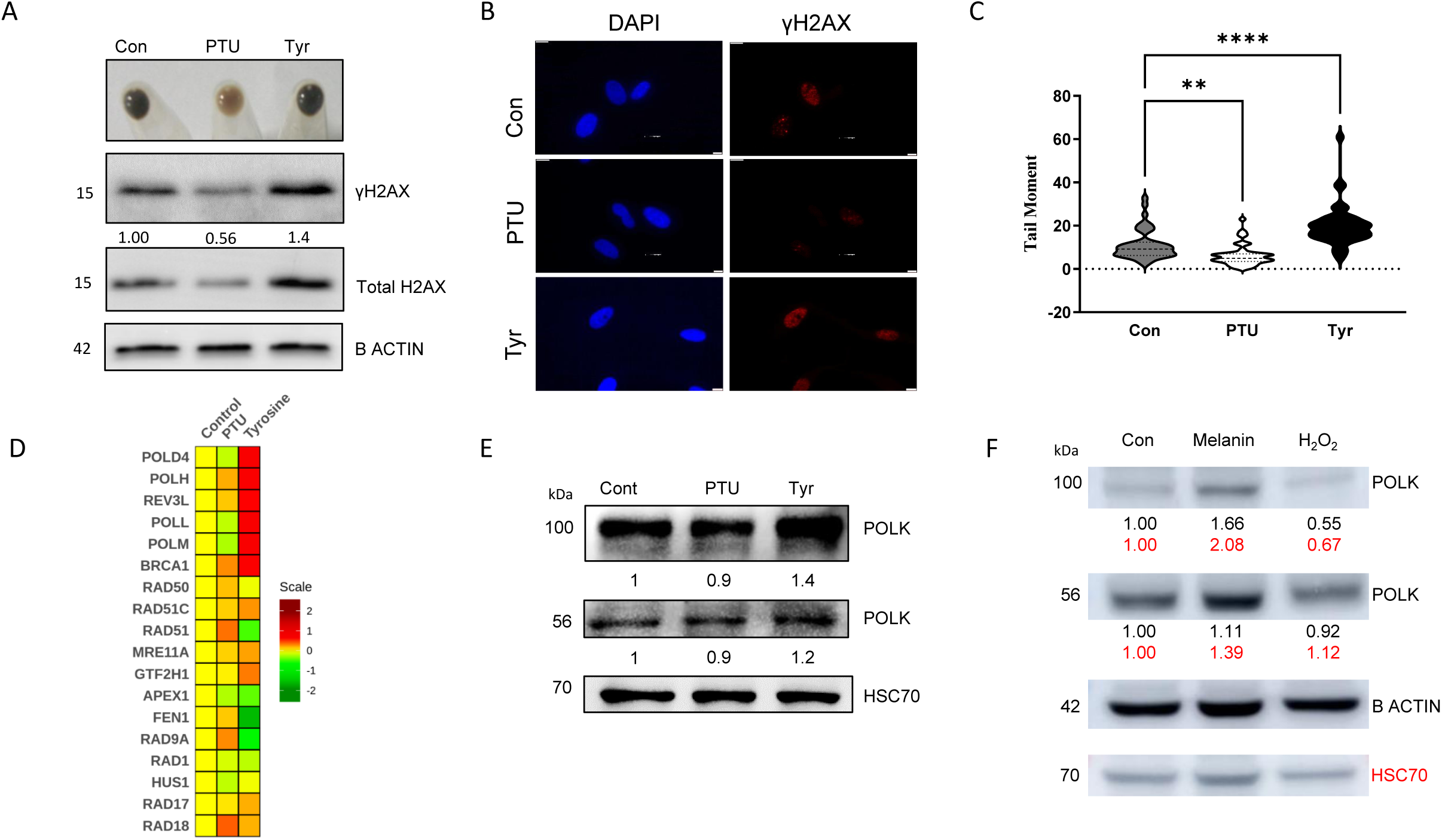
Normal Human Epidermal Melanocytes (NHEM) respond to pigmentation induced DNA damage by elevating PolK. A. NHEM cells were treated with 200 µM PTU or 1mM Tyr for 5 days for differential pigmentation. (Top) Cell pellet, (bottom) western blot analysis of cell lysates with phosphorylated H2AX, total H2AX and beta Actin antibodies. Numbers represent relative fold change wrt beta actin. B. Immunofluorescence of NHEM treated with PTU and Tyr with phosphorylated H2AX antibody that labels double strand DNA breaks. C. PTU and Tyr treated NHEM cells were subjected to single cell electrophoresis and comet analysis. Mean tail moment distribution across each population of duplicate biological experiments with atleast 50 comets analyzed is depicted by a violin plot. Error bars represent ± SEM. students unpaired t test ns non-significant, *** p val < 0.001, **** p val< 0.0001 D. Gene expression by microarray analysis of NHEM cells treated with PTU or Tyr for 5 days for the set of DNA repair genes is depicted as a heat map. E. Western blot analysis of PTU and Tyr treated cells with PolK antibody and normalized to Hsc70. Numbers indicate normalized fold change in PolK level wrt Hsc70. F. Western blot analysis of B16 cells transfected with only DNA (con) or melanin-modified DNA or oxidized by H2O2 with Pol κ antibody normalized to beta-actin and HSC70.

Subjecting primary melanocytes with varying levels of pigmentation to microarray analysis identified several DNA repair genes to be differentially expressed (**Fig 4D**) (Motiani et al., 2018). Validating the B16 based observations, several of the replication stress response genes were elevated in primary melanocytes. Of these, Pol κ at the protein level was induced upon hyperpigmentation and reduced upon depigmentation (**Fig 4E**). Hence the response of primary human melanocytes to pigmentation appears to be the induction of this translesion polymerase enzyme possibly to achieve reversal of DNA damage caused by melanin. Extraneous transfection of plasmid DNA modified with intermediates of melanin but not oxidized by H_2_O_2_ in depigmented B16 cells further substantiated this possibility (**Fig 4F** & **Fig S2D**). We then proceeded to investigate the role played by this repair system in maintaining genome integrity and cellular homeostasis.

### Silencing of Pol κ abrogates pigmentation induced replication stress

CRISPR-based genetic ablation of this polymerase could possibly result in multiple mutations. Hence, we resorted to silencing Pol κ in B16 cells stably expressing shRNA targeting Pol κ and this was compared to non-targeting shRNA against luciferase. These cells were freshly transfected and used as an enriched pool to prevent confounding effects of translesion repair alterations possibly affecting genome integrity. When these cells were allowed to pigment, comparable pigmentation could be achieved by day 7 (**Fig 4B** top). Interestingly, the number of cells were significantly higher upon Pol κ knockdown. Cell cycle analysis revealed an increase in S+G2/M population suggesting lack of cell cycle arrest despite pigmentation (**Fig 5A** bottom).

**Figure 5:**
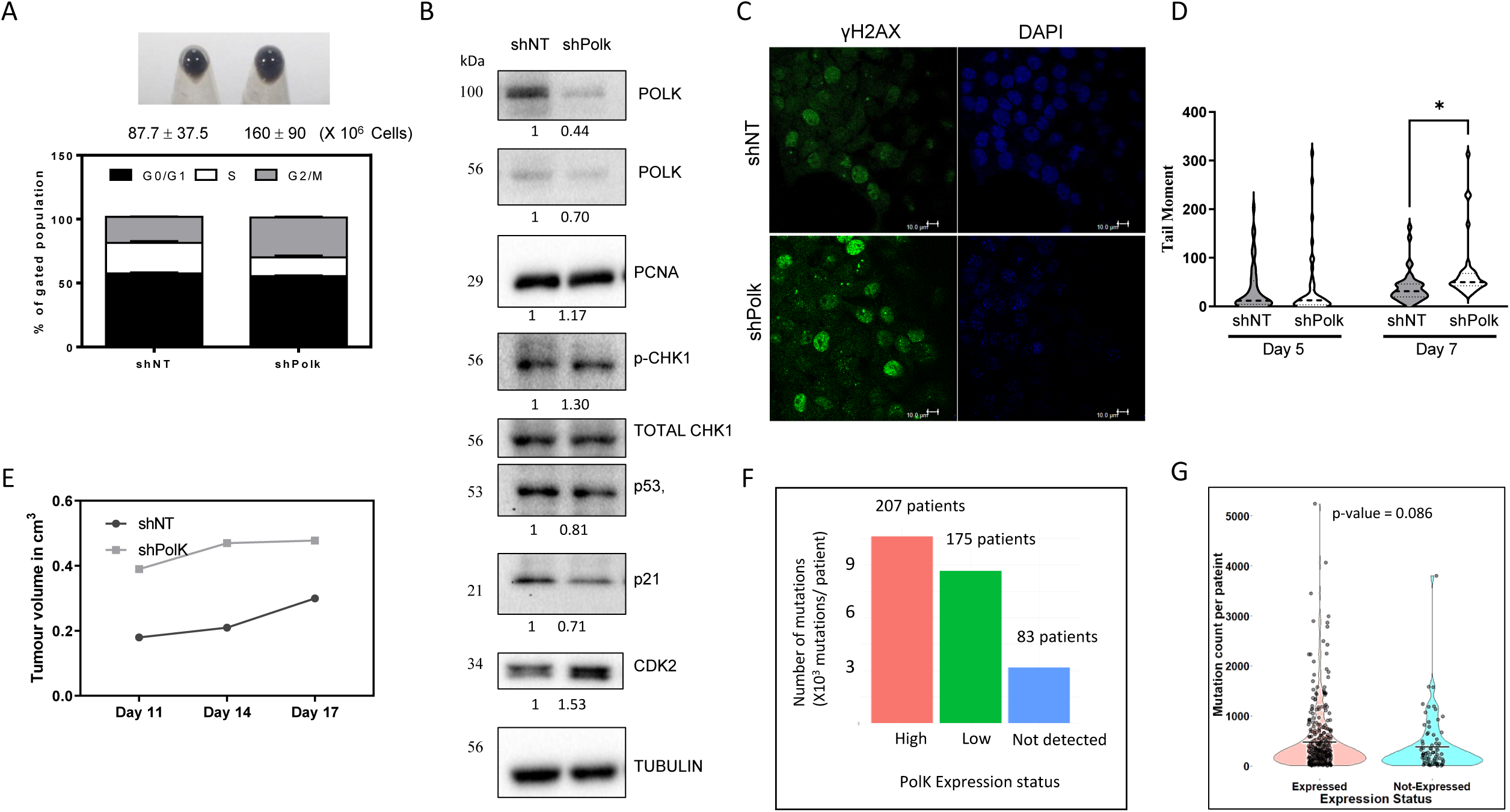
During pigmentation silencing of Polk prevents cell cycle arrest, despite elevated DNA damage. A. (top) Cell pellets of control non-targetting (shNT) and *Polk* silenced (shPolk) B16 cells on day 7 of pigmentation. Numbers below represent mean ± SEM cell counts across biological triplicates. (bottom) Cell cycle analysis stable B16 cells. B. Westernblot analysis of DNA repair and cell cycle related proteins in control and shPolk cells. Numbers below represent tubulin normalized fold changes. C. Immunofluorescence analysis of control and shPolk cells with phosphorylated H2AX antibody that labels double strand DNA breaks (puncta labelled in green) and the nucleus is counterstained with DAPI. D. Control (shNT) and shPolk expressing pigmented B16 cells at day 7 were subjected to single cell electrophoresis and comet analysis. Mean tail moment distribution across each population of duplicate biological experiments with atleast 50 comets analyzed is depicted by a violin plot. Error bars represent ± SEM. ns non-significant, **** p val < 0.0001 E. Control (shNT) and shPolk expressing B16 cells were injected inside the flank of C57/BL6 mice and allowed to grow as tumors. The volume of the tumor was non-invasively monitored and plotted over time. F. Analysis of melanoma samples from TCGA data for mRNA expression of *Polk* and mutation burden per sample. 83 samples were binned based on the expression of *Polk*. Mutation burden in each group is depicted as a bar plot. G. Mutational count per patient across *Polk* expressing (high+low) and not expressing samples is depicted. Statistics employed Welch two-sample t-test. P-value < 0.08.

A panel of replication check point control orchestrators and effectors were analyzed by western blot on day 7 of pigmentation. Confirming the silencing, levels of both 100 as well as 56 kDa forms of the Pol κ protein were downregulated (**Fig 5B**). PCNA is an effector of Pol κ and a marker of proliferating cells. However, upon Pol κ silencing PCNA levels remained unaltered, perhaps due to opposing effects of damage and proliferation in the absence of key repair polymerase. Interestingly, p-Chk1 levels were elevated suggesting that changes upstream of sensing of DNA damage is operational. However, both p53 and p21 that function to couple DNA damage to cell cycle arrest were low and Cdk2 was elevated. Thereby more cells are in the proliferative phase of cell cycle.

### Silencing of Pol κ results in proliferative melanocytes with damaged DNA

Abrogation of cell cycle arrest could be due to resolution of DNA damage. However, Pol κ silenced cells showed higher γH2AX foci (**Fig 5C**). Analysis of DNA damage by comet indicated that the two cell types had comparable and low levels of damage during early stages of pigmentation, however on day 7 the Pol κ silenced cells had significantly higher DNA breaks suggesting that the DNA repair kinetics is slower in the absence of this translesion polymerase (**Fig 5D**). We confirmed that transient silencing of Pol κ between day 5 to day 7 of pigmentation using a sequence independent siRNA resulted in a similar increase in tail moment (**Fig S5A** & **B**). Therefore, we confirmed that the observations made with the shRNA clone could be recapitulated by the sequence independent siRNA pools. We resorted to the shRNA-based comparisons for long term experiments. An increase in the abasic sites was observed the Day 7 Pol κ silenced cells which is in accordance with the increased comets observed earlier (**Fig S5C**). Hence the observed lack of cell cycle arrest coupled to higher damage is indicative of altered repair check point activation, and the cells are likely to accumulate higher mutational burden. Upon injection of shNT control and shPol κ cells into the flank of C57/BL6 mice to grow autologous pigmented tumors, consistently shPol κ cells formed bigger and aggressive pigmented tumors (**Fig S5D, S5E** & **Fig 5E**). Thereby confirming that Pol κ is a critical response mounted by melanocytes to tackle melanogenesis induced DNA damage, and in its absence the critical cell cycle checkpoint is circumvented, potentially rendering the cells vulnerable to genome instability. Substantiating this possibility, a recent systematic investigation of combination of DNA damaging agents and repair gene knockouts carried out in *C. elegans* highlighted a heightened mutational frequency for alkylating agents MMS and DMS upon deletion of the Pol κ ortholog (Volkova et al., 2020).

### Signatures of somatic mutations in human melanoma with varying Pol κ levels

Having established the role of melanin as a DNA damaging agent and mapping the error-prone response by melanocytes, we decided to explore whether this is relevant in the human melanoma context. We extracted the cutaneous melanoma samples from The Cancer Genome Atlas (TCGA) and segregated the samples based on the mRNA expression of POLK (Weinstein et al., 2013). While 83 melanoma samples did not have detectable expression of POLK mRNA by sequencing, FPKM based binning resulted in 207 patients with high levels of POLK and 175 patients had lower expression of POLK (Uhlen et al., 2017). Overall number of somatic mutations were highest in high-POLK group and least in samples that did not have detectable POLK. A systematic two group comparison of somatic mutation burden using Welch Two Sample t-test in POLK expressing (high+low) with POLK non-detected group resulted in a p value < 0.08 and the mean in group “expressed” is 505 whereas the mean in group “not-expressed” is 380. Thereby, suggesting that repair by Pol κ could explain a fraction of mutations in human melanoma, and provide physiological relevance to the observations in cultured cells. This provides a compelling reason to further investigate this aspect of eumelanogenesis and its implication in melanoma.

## Discussion

While most non-primate mammals employ pigmentation of hair, humans additionally sport a pigmented skin. In evolutionary timelines, acquisition of epidermal pigmentation is a recent innovation among humans and has provided the much-needed protection of the naked skin from UV rays. This adaptive feature has enabled human colonization in several latitudes that differ in the incident UV radiations, and hence is believed to be beneficial. While the UV protection role is likely to be overarching, the current work unravels a rather unwanted side-effect during the synthesis of this esoteric biopolymer. In this work, we demonstrate that uncontrolled melanin synthesis causes DNA damage. The key response of pigmenting cells is the translesion repair by Pol κ via the ATR-Chk1 signaling axis. When Pol κ levels fail to rise during the surging pigmentation response such as recurrent tanning, melanocytes are unable to elicit checkpoint activation and could result in accumulation of mutations.

The known role of Pol κ in repairing benzo[a]pyrene adducts, as well as alkylating agents such as MMS modification strongly supports analogous role for repairing melanin-adducts with DNA (Takenaka et al., 2006; Zhang et al., 2000). Since melanin is a biochemically complex and insufficiently characterized polymer, formal demonstration of covalent adduct formation using analytical methods is extremely challenging. Among translesion repair enzymes, Pol κ is necessary to mend large bulky adducts on the nucleotide bases that are not amenable for repair by other DNA repair systems (Bi et al., 2005). *In vitro* experiments highlight this promiscuity but have raised the question on the *in vivo* fidelity of repair, which has been difficult to assess (Ohashi et al., 2000). The recent work by Volkova NV et., al systematically investigated multiple combinations of DNA damaging agents with genetic mutants of different repair components in *C. elegans*. In this study Pol κ emerged as a prime candidate for MMS and EMS induced DNA repair. The authors report several-fold increase in base substitution frequency observed in Pol κ KO, that indicates repair of alkylating agents by Pol κ. As the knock-out strategy underscores the repair capacity of the cell in the absence of Pol κ, based on our study, we hypothesize that absence of replication stress response could contribute to the observed genome instability.

Elevated expression as well as mutations in Pol κ gene is observed in several cancers (Bavoux et al., 2005; O-Wang et al., 2001; Peng et al., 2016; Wang et al., 2010). In an earlier study authors report subcellular localization of Pol κ inside nucleus of the human melanoma cells and this involves mTOR pathway (Temprine et al., 2020). The work further demonstrated Pol κ to provide survival advantage upon pharmacological inhibition of the oncogenic BRAF, independent of creating somatic mutations in the genome. In this study overexpression of *PolK* did not significantly increase the mutational burden, it is likely that the Pol κ-based repair of melanin modified DNA is indeed error -prone but in the absence of this critical repair system the associated errors could be several fold higher. In another study role of Pol κ in promoting DNA synthesis and recovery of replication stress at stalled forks upon nucleotide deprivation was elucidated (Tonzi et al., 2018). Cells seems to have multiple regulatory check points at the level of *Polk* gene expression as well as nuclear localization of Pol κ, possibly due to its error-prone nature of repair and/or its crucial role in replication fork recovery.

Most of the work on UV-induced melanoma formation have concentrated on direct DNA damage. This is substantiated by the prevalence of UV-signature mutations in several melanoma samples. Additionally, intermittent exposure to UV is also known to initiate melanomagenesis by involving the immune effector system through chemokines which are believed to promote this form of cancer (Zaidi et al., 2011). These studies are supported by demographic data on the incidence of melanoma among people of European origin in Australia. These individuals inherently have a low melanin content of skin but are exposed to higher UV doses during sun-tanning and are highly predisposed to melanoma formation. Recent data suggests an increase in the incidence of melanoma among users of artificial tanning beds (Ghiasvand et al., 2017). Surprisingly, despite deep-seated within the epidermis and loaded with melanin, melanocytes are not well-protected from UV and are vulnerable to DNA damage.

In melanoma, it is apparent that there are both UV-signature mutations and unexplained large-scale genomic alterations associated with malignant cells that are thought to be passenger mutations. Specifically, the known key oncogenic mutations in melanoma including BRAF V600E and NRAS Q61L/R do not have the UV-signature (Hodis et al., 2012). It is likely that UV-induced melanogenesis could result in these driver mutations adding a layer of complexity to the link between UV and melanoma. While the activation of melanocytes in conditions such as tanning enable adaptive pigmentation response, repeated or uncontrolled activations could alter these cells and possibly promote mutagenesis and facilitate their transformation into melanoma. Thereby our study establishes a hitherto unknown, undesirable side of melanin and its possible involvement in melanoma.

## Acknowledgement

Council for Scientific and Industrial Research (CSIR), India, through grant RegenX (MLP2008) and Department of Biotechnology through the grant (GAP0182) provided support to the execution. SK and MG acknowledge CSIR for Research Fellowship. We thank Dr Vamsi K Yenamandra for the meaningful discussions and constant support throughout the execution of this work.

## Materials and Methods

### Cell lines, Reagents and Media

B16 cells were obtained from ATCC. Primary human melanocytes (Normal Human Epidermal Melanocytes, NHEM) were obtained from Lonza. All cell culture reagents were obtained from GIBCO Life technologies. siRNAs and shRNAwere purchased from Dharmacon (GE Healthcare) and transfections were performed using Dharmafect II for siRNAs and Lipofectamine 2000 for shRNAs. Midiprep plasmid preparation was performed using Qiagen Midi DNA kit. KAPA SYBR FAST qPCR Master Mix was obtained from KAPA Biosystems. Genomic DNA and RNA isolations were performed using Macherey Nagel NucleoSpin® TriPrep kit (Cat no: 740966).

### Setting of low density (LD) Pigmentation model of B16 mouse melanoma cells

Cultures were set up as described by (Natarajan et al., 2014). Briefly, depigmened B16 cells were seeded at a density of 100 cells/cm^2^ in DMEM high glucose media (Sigma; D5648) with 10% FBS (Invitrogen, 10270) to make them pigmented by day 6 to day 8. During the span of pigmentation the media was not replenished.

### Culturing and chemical treatments on melanocytes

B16 were cultured in DMEM high glucose and 10% FBS, and primary human melanocytes were grown in M254 (Gibco, M-254-500), both at 5% CO_2_ concentration. All treatments were given at 24h post seeding. 200uM of Phenylthiourea (PTU, sigma; P7629) was used was depigmentation, and 1mM of L-Tyrosine was used for hyperpigmentation of cells.

2mM of Hydroxyurea (sigma; H8627) for 3h was used for inducing replication stress, 5 J/m^2^ of UVA and 200mJ/cm^2^ of UVB were used as photooxidative damaging agents. 100mM H_2_O_2_ for 15 min was used as oxidative stress inducer. AZ20 (Sigma; SML1328) was used as ATR inhibitor at 50nM concentration.

### Detection of DNA base modifications by ELISAs

Genomic DNA was isolated using Macherey Nagel Nucleospin Triprep kit. Abasic sites and 8-oxoGuanine were determined using ELISA kits from cell biolabs (OxiSelect™ Oxidative DNA Damage Quantitation Kit, AP sites, STA-324 and OxiSelect™ Oxidative DNA Damage Quantitation Kit, 8-OHdG, STA-320) using manufacturer’s instructions.

### Detection of Reactive Oxygen Species and Apoptotic cells

Relative levels of reactive oxygen species were determined using CM-H2DCFDA(2′-7′-Dichlorodihydrofluorescein diacetate) assay kit (Thermo, Cat no.C6827). Detection of apoptotic cells was done using The Alexa Fluor® 488 annexin V/Dead Cell Apoptosis Kit (Invitrogen, V13241) as per manufacturer’s protocol.

### Cell cycle analysis of B16 cells

B16 cells were trypsinised and washed once with PBS. Cell pellet of approximately 3-5 million cells was resuspended in 200-300 µl of 70% chilled ethanol and was stored overnight in minus 20^0^C. For analysis, cells were spun at 7000 rpm for 10 minutes at 4^0^C. Supernatant ethanol was discarded and washed thrice with 1X PBS to remove traces of ethanol. Finally, cells were resuspended in 100-200μl of (PBS+0.1%Triton X100) along with RNAase A (sigma; R6513) to a working concentration of 0.5mg/ml. These were incubated overnight at room temperature. After taking out some cells as unstained control, 2μl of propidium iodide (2mg/ml) was added to approximately 100μl of cell suspension and incubated at room temperature for 10 minutes in dark. Cells were then directly used for flow cytometric analysis.

### Cell fractionation and western blot analysis

For whole cell lysate B16 cell pellet was resuspended in NP40 Lysis Buffer (Thermo; ALF-J60766-AP) reconstituted with protease and phosphatase inhibitors (750μl of NP40 lysis buffer, 1X PIC, 10mM sodium pyrophosphate, 1mM sodium orthovandate, 10mM sodium glycerophosphate, 1mM PMSF). Cells were incubated in lysis buffer overnight at -80^0^C and centrifuged at 13000 rpm for 20 minutes at 4^0^C, protein supernatant was collected and estimated using standard BCA protocol (Pierce BCA protein assay kit; Thermo). Cellular fractionation was performed by resuspending the cell pellet in hypotonic Buffer A (10 mM HEPES, pH 7.9, 10mM KCl, 1.5mM MgCl_2_, and 0.5mM DTT) while keeping the pellet on ice and incubating for 5 min. Cells were further lysed using strokes of a hand-held homogenizer. When more than 70% of cellular lysis was achieved as visualized under 40X microscope, suspension was centrifuged at 1000 rpm for 5 minutes at 4^0^C. Supernatant (cytosolic fraction) was separated and nuclear pellet was processed similar to whole cell pellet. 30μg of the protein was used for proteins in the size range of 20-150 kDa. Primary antibody incubations were done for overnight in cold room (4^0^C). Subsequent to incubation with the primary antibody, the membrane was washed thrice with TBST containing 0.1% tween and were further incubated with corresponding HRP conjugated secondary antibody at 1:5000 dilution in 5% BSA 5% Skim milk or for one hour at room temperature. Membrane was washed in TBST with 0.1% Tween-20, thrice for 15 minutes each and developed using ECL reagent.

### Antibodies

Following Primary antibodies were used. Pol κ (ab57070), γH2AX(CST 9718), total H2AX(CST 2595), pCHK1 (CST 2348P), Total CHK1 (Invitrogen PA512096), pATR (CST 2853), total ATR(CST 2790), p53(ab38497), p21(CST 9706), PCNA (SC56.), DCT (ab74073) and TYR (custom synthesized genscript). Secondary antibodies conjugated to Alexa fluor for immunofluorescence were procured from Life technologies (USA). Secondary antibodies conjugated to HRP required for western blot were obtained from GE Healthcare.

### Immunocytochemistry on B16 cells

Cells were seeded on UV treated sterile coverslips (Corning) in six well plates. Cells were fixed with 2% paraformaldehyde in PBS for 10 minutes at 37^0^C. Further, they were permeabilized with 0.1 % Triton® X-100 in PBS for 15 minutes at room temperature on slow orbital shaking. After three consecutive washes with 1X PBS, cells were blocked with 5 % NGS (Jackson’s immunoresearch) for 1 hr at room temperature on slow orbital shaking. Cell were given a single wash of PBS and incubated with γ H2AX antibody (CST 9718, 1:200) or anti Pol κ (ab97070, 1:100 in 1%NGS) for two hours at room temperature. Post primary antibody incubation cells were washed thrice with PBST (PBS+0.1%TritonX 100) for 5 minutes each on slow orbital rotation and then incubated in secondary Alexa Flour 568 anti-mouse antibody (1:500 in

1%NGS) for 1 hr at room temperature in dark. Finally, cells were washed thrice with IX PBST and mounted on glass slides with 10μl of antifade DAPI and sealed the coverslips using acetone. Imaging was done using Leica confocal microscopy and quantitated using LAS software (Leica). Average nuclear fluorescence intensity was determined using Leica Application suite AF (LAS AF) software.

### Generation of Tyrosinase mutant B16 cells

Tyr CRISPR was designed using ECRISP (http://www.e-crisp.org/E-CRISP/) After overlap PCR of the CRISPR, it was in vitro transcribed using mMessage mMachine T7 ULTRA kit (Thermo; AM1345). For transfections, B16 melanoma cells were seeded at a regular density of 2 lakh cells per well of a six well plate. A complex of CRISPR RNA with Cas 9 protein was prepared and incubated for 20 minutes at room temperature. 500μl of optiMEM (gibco; 31985070) along with 3μl of Lipofectamine 2000 (Invitrogen; 11668019) was added to the above mix and incubated again for 20 minutes. After incubation, media was removed from culture and given a single was of 1X DPBS. CRISPR mix was added to these cells after making up the volume to 1ml with fresh optiMEM. This was incubated for 6 hrs after which media was changed to terminate transfections. These transfected cells were used for setting up multiple LD culture flask in T75. The depigmented colonies were manually picked up by visual selection under microscope followed by *in situ* trypsinization on Day7 of LD culture. These colonies were expanded and analyzed for tyrosinase levels and activity.

### siRNA transfections in B16 melanoma cells

B16 cells were seeded in 6-well plates at a density of 2 × 10^5^ cells per well. Transfections were carried out with Dharmafect II reagent as per manufacture’s protocol. After 48 hours the cells were trypsinised and pelleted for RNA or protein isolations. For low density cultures (LD), cells were transfected on Day 3 of cycle and terminated on Day 6. During transfections, culture media was stored in sterile falcon or flask. After transfections, cultures were given a single wash of DPBS and replaced back with same media for cells to normally pigment during the course of LD cycle.

### shRNA transfections and stable knockdown of *Polk* in B16 melanoma cells

All shRNA’s (GIPZ Lentiviral constructs) were purchased from Dharmacon (GE Healthcare). Pol κ shRNA (RMM4532-EG27015) glycerol stocks were used for generation of stable cell lines for Pol κ knockdown. B16 cells were transfected with *Polk* silencing plasmid. 24h post transfections, selection of stable cells was done by treatment with Puromycin. Cells were kept under selection pressure for 3-4 weeks by regular change in media with fresh addition of puromycin every alternate day. In case of confluency, cultures were passaged, with treatment starting at 24 h post seeding. This was done until all the cells showed consistent GFP fluorescence.

### *In vivo* model of melanoma induction in mice

All animal procedures were done in accordance with the Indian Committee for the Purpose of Control and Supervision of Experiments on Animals (CPCSEA), (Section 15(1) of the Prevention of Cruelty to Animals Act, 1960; Registration number.38/GO/Re Bi/SL/99/CPCSEA). **Institutional Animal Ethics Committee, under Indian Association for the Cultivation of Science (IACS)** approved all experiments and procedures carried out on the animals. For induction of melanoma tumors, Pol κ knockdown stables or the non-silencing control cells were injected 5 outbred male/female C57BL/6 mice each at 4–6 weeks of age (National Institute of Immunology, Delhi). A single subcutaneous administration of 10^6^ cells in 100μl volume of cell suspension in right hand side flank region was given. The time and appearance of the first tumor (latency period) as well as the number of mice with tumors (incidence) was recorded during the study. At different time intervals i.e., on Day 11, Day 14 and Day 17, tumour was excised; length and width were measured using a Vernier Caliper.

### Statistical analysis and graphs

GraphPad prism was used to plot the graphs. Statistical analysis to obtain significance in the data was performed using Student’s *t*-test. P value(P) > 0.05 is marked as ns ≤ 0.05 is marked as *, P ≤ 0.01 as **, P ≤ 0.001 as ***and P ≤ 0.0001 as ****.

## Supplementary Figure Legends

**Supplementary Figure S1:**

**Exploring the effect of melanogenesis on DNA damage and elicitation of repair pathways**

A. Schematic of the experimental design employed in the study to alter pigmentation of melanocytes by the use of tyrosinase inhibitor Phenylthiourea (PTU) and its substrate L-tyrosine (Tyr). (Right) coupling of differential pigmentation to temporal resolution to dissect the pigmentation DNA repair interplay.

B. Estimation of early apoptotic (annexin V positive population), late apoptotic (annexin V and PI double positive population) and dead (sub G0 population) upon differential pigmentation. Bars represent mean ± SEM across two biological replicates.

C. Cell cycle analysis of high density (non-permissive for pigmentation) B16 cells at 24h and 48h post treatment with PTU and Tyr. Stacked bars represent mean ± SEM across two biological replicates.

D. Cell cycle analysis of B16 cells at day 5 (early pigmentation) day 6 (mid-phase of pigmentation) and day 7 (late phase of pigmentation). Stacked bars represent mean ± SEM across two biological replicates.

E. Cellular ROS content was assayed with 2’,7’–dichlorofluorescin diacetate (DCFDA) staining and mean fluorescence value across duplicate biological experiment is depicted as a bar plot. Bars represent mean ± SEM. students unpaired t test * p val< 0.05

F. ELISA based 8-OHdG estimation in control, PTU and tyrosine treated cells on day 7 of pigmentation induction.

**Supplementary Figure S2:**

**Melanogenesis alters DNA, induces DNA damage and elicits expression of translesion polymerase *Polk***

A. Western blot analysis of control, PTU or tyrosine treated day 7 treated B16 cells with Pol κ antibody. Numbers represent relative fold change *wrt* beta actin.

B. PTU, Tyr and both treated day 7 pigmented B16 cells were subjected to single cell electrophoresis and comet analysis, representative images of PI-stained comets are depicted.

C. Microarray analysis of B16 cells on different days of pigmentation. Fold change is represented as a heat map *wrt* Day 0. Two-way hierarchical clustering of genes based on the expression pattern is depicted.

D. Agarose gel electrophoretic mobility assessment upon treatment of pLH-sgRNA-Sirius-8XPP7 plasmid DNA with ongoing non-enzymatic gradual melanogenesis (lane 2), enzymatic melanogenesis (lane 3) and treatment of DNA with pre-synthesized melanin (lane 4).

**Supplementary Figure S3:**

***Polk* is elicited as a key DNA translesion repair component upon melanogenesis induced DNA damage**

A. Real-time qRT-PCR analysis of select homologous recombination-based DNA repair genes during different days of pigmentation in B16 cells.

B. Real-time qRT-PCR analysis of select translesion DNA repair genes during different days of pigmentation in B16 cells.

C. Western blot analysis of B16 cells at different days of pigmentation Pol κ and beta Actin antibodies. Numbers represent relative fold change wrt beta actin.

D. Quantitation of mean nuclear fluorescence intensity of Pol κ of differentially pigmented day 7 B16 cells treated with PTU, Tyr or both (image depicted in Fig 2C.). Bars represent mean ± SEM across two biological replicates.

E. Real-time qRT-PCR analysis of *Polh* translesion polymerase in differentially pigmented B16 cells treated with PTU and Tyr.

F. Real-time qRT-PCR analysis of *Rev3l* translesion polymerase in differentially pigmented B16 cells treated with PTU and Tyr.

**Supplementary Figure S4:**

**Targeted ablation of Tyrosinase locus with CRISPR to generate a genetic model of hypopigmentation**

A. The mouse Tyrosinase (*tyr*) Gene architecture and the position of single guide RNA sequence.

B. Sanger sequencing chromatogram of wild type and *tyr* mutant clones. Deletion of C nucleotide at position 459 is indicated by the blue arrow.

C. Predicted amino acid sequence of WT and Tyr^fs118^ coding region.

D. Western blot analysis of B16 WT and Tyr^fs118^ cells on day 5, 6 and 7 post induction of pigmentation.

Numbers below represent WT day 5 fold normalized expression wrt *Gapdh*.

E. Confocal images of B16 WT and Tyrosinase mutant (Tyr^fs118^) at Day 7. Newly synthesised DNA was labelled using EdU analog and tagged by click chemistry (in red). Immunofluorescence using γH2AX antibody is labelled (in green). Bright field images indicate the presence of melanin granules. Scale bars represent 10um.

F. Quantitation of mean EdU nuclear intensity across 50 cells is depicted as a bar representing mean ± SEM. students upaired t test * p val< 0.0001

G. Quantitation of mean γH2AX nuclear intensity across 50 cells is depicted as a bar representing mean ± SEM. students upaired t test * p val< 0.0001

**Supplementary Figure S5:**

**Silencing of *Polk* by an siRNA pool, increases DNA damage and B16 clone stable expression of shPolk results in increased melanoma growth**

A. Polk was silenced using an siRNA pool and realtime PCR analysis of the *Polk* mRNA levels were measured by qRT-PCR analysis.

B. B16 cells transfected with control non-targeting or *Polk* silencing siRNA pool on Day 3 of pigmentation and on day 5 subjected to single cell electrophoresis and comet analysis. Mean tail moment distribution across each population of duplicate biological experiments with atleast 50 comets analyzed is depicted by a violin plot. Error bars represent ± SEM. Student’s t-test (unpaired), ns non-significant, ** p val < 0.01

C. Number of abasic sites in the genomic DNA of B16 cells stably expressing non-targeting shRNA or *Polk* silencing shRNA. Bars represent mean ±SEM across duplicate biological experiments. Student’s unpaired t test **** p val < 0.0001

D. Mice images on day 17 post-injection of control (shNT) and shPolk expressing B16 cells into the flank of C57/BL6 mice and allowed to grow as tumors.

E. Images of excised tumours of control and *Polk* silenced B16 cells injected in C57/BL6 mice.

